# Discovery of small molecule inhibitors of Plasmodium falciparum apicoplast DNA polymerase

**DOI:** 10.1101/2022.03.04.482688

**Authors:** Supreet Kaur, Nicholas Nieto, Peter McDonald, Josh R. Beck, Richard B. Honzatko, Anuradha Roy, Scott W Nelson

**Author notes:** **Disclosure statement.** The authors report there are no competing interests to declare.

## Abstract

Malaria is caused by infection with protozoan parasites of the *Plasmodium* genus, which is part of the phylum Apicomplexa. Most organisms in this phylum contain a relic plastid called the apicoplast. The apicoplast genome is replicated by a single DNA polymerase (apPOL), which is an attractive target for anti-malarial drugs. We screened small-molecule libraries (206,504 compounds) using a fluorescence-based high-throughput DNA polymerase assay. Dose/response analysis and counter-screening identified 186 specific apPOL inhibitors. Toxicity screening against human HepaRG human cells removed 84 compounds and the remaining were subjected to parasite killing assays using chloroquine resistant *P. falciparum* parasites. Nine compounds were potent inhibitors of parasite growth and may serve as lead compounds in efforts to discover novel malaria drugs.

## Introduction

Malaria kills over half a million people each year with the majority of those deaths occurring in children under five years of age ^1^. Over 40% of the world’s population lives in areas where Malaria is a serious health risk with more than 215 million new cases diagnosed each year^1^. During the last century, Malaria was controlled with chloroquine, but resistance emerged in the 1950’s, starting in Southeast Asia, then spreading throughout Asia, and finally Africa ^2^. Artemisinin and its derivatives are current drugs of choice and are especially effective in combination with slow-acting anti-malarial drugs. In 2005, partial resistance to artemisinin arose in parasites from Cambodia, Myanmar, Thailand and Viet Nam ^3^. Parasites resistant to artemisinin-based therapies are now prevalent in Southeast Asia and are emerging in Africa, jeopardizing global programs to control Malaria. New drugs that targeting novel parasite biology may play a critical role in curbing the impact of this disease ^4-7^.

Malaria is caused by parasites of the genus *Plasmodium* of the phylum Apicomplexa ^8^. Nearly all organisms within this phylum contain an unusual organelle called the apicoplast. The apicoplast is evolutionarily related to the chloroplast and participates in several metabolic pathways, including the biosynthesis of fatty acids, heme, iron-sulfur clusters, and isoprenoids ^9^. The parasite relies on the apicoplast for the synthesis of isoprenoids. Defects in apicoplast metabolism, or its failure to replicate and divide, leads to the death of *Plasmodium* at the blood and liver stages of infection ^10^. Hence, the apicoplast is a promising drug target ^11^.

Nearly 600 proteins exist within the apicoplast and approximately 4% of those are involved in replication of the 35 kb apicoplast genome ^12^. Enzymes that carry-out the fundamental process of DNA replication are promising drug targets. Indeed, ciprofloxacin, a DNA type II topoisomerase inhibitor, has confirmed anti-malarial activity; however, antiparasitic properties could be due to off-target effects instead of, or as well as, inhibition of the *P. falciparum* gyrase ^13^ The apicoplast DNA polymerase (apPOL) represents an attractive target for anti-malarial drug development ^14^. apPOL exhibits low amino-acid sequence identity to replicative polymerases from other species. Outside of Apicomplexa, the nearest homolog (35% sequence identity) is the replicative polymerase from the cyanobacteria Cyanothece sp. PCC 8802. The most similar human DNA polymerases are the lesion bypass polymerases theta and nu (23% and 22% sequence identity, respectively), with the other human DNA polymerases displaying less than 20% identity, providing a foundation for selectivity. On the other hand, apPOLs from the two primary causative agents of human Malaria (*P. falciparum* and *P. vivax*) are 84% sequence identical, suggesting that drugs targeting *P. falciparum* apPOL would be effective in treating Malaria caused by *P. vivax*.

To discover novel inhibitors of *P. falciparum* apPOL (hereafter, *Pf*-apPOL), we screened of several commercial small-molecule libraries using a fluorescence-based high-throughput DNA polymerase assay. The primary hits were subjected to dose/response analysis and counter-screening against bacteriophage T4 DNA polymerase to remove non-specific inhibitors and compounds that interfered with the assay. The specific *Pf*-apPOL inhibitors (186) were screened for toxicity against human cells and non-toxic compounds (102) were tested in parasite killing assays using chloroquine resistant *P. falciparum* parasites cultured in human red blood cells. Ultimately, nine inhibitors passed all screens and arrested parasite growth in human red blood cells.

## Materials and methods

### Materials and reagents

Codon-optimized, exonuclease-deficient *Pf*-apPOL was overexpressed in *E. coli* BL21 (DE3) and purified as described ^15^. The protein was dialyzed against two, 1-liter changes of 20 mM Tris-HCl (pH 8.0), 400 mM NaCl, and 20% glycerol. *Plasmodium bergei* and *Babesia bovis* apPOLs (*Pb*-apPOL and *Bb*-apPOL, respectively) were codon-optimized, expressed, and purified as for *Pf*-apPOL^15^. Bacteriophage T4 DNA polymerase was purified as described ^16^. The DNA oligonucleotides used to make the high-throughput substrate were obtained from Integrated DNA Technologies. Deoxynucleotides were purchased from Strategene. All other materials came from Fisher Scientific.

### High-throughput (HT) Assay

The HT assay employs a molecular beacon, a stem-loop DNA template with a Cy3 dye and the Black Hole Quencher on the 3′ and 5′ ends of the DNA, respectively ^15^. *Pf*-apPOL concentration is based on a calculated absorptivity of ε280 = 56750 M^-1^ cm^-1^. Reactions were carried out at 25°C, using Corning 3575 386-well plates (well-volume, 20 μL). Assays testing for compound inhibition composed of reaction buffer (pH 7.9 in 50 mM potassium acetate, 20 mM Tris-acetate and 10 mM magnesium acetate), *Pf*-apPOL (20 nM), DNA substrate (20 nM), dNTPs (6 μM), and compound (10 μM in DMSO). The DNA and nucleotide substrates (dNTPs) were at concentrations of 20 nM and 6 μM, respectively, in the assay. *Pf*-apPOL was incubated in the reaction buffer with dNTPs and the compound of interest (10 μM, in the assay) at 25°C for 10 minutes. The DNA substrate was then added and the reaction was allowed to proceed for 90 minutes at 25°C.

Each assay plate contained 16 negative controls (no *Pf*-apPOL, DMSO present) and 16 positive controls (*Pf*-apPOL present, no DMSO). The fluorescence of each well was determined using a BioTek Synergy Neo fluorescence plate reader using excitation and emission wavelengths of 555 and 570 nm, respectively. The % inhibition was determined using equation 1:

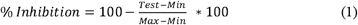

In the above, *Min* is the average of fluorescence emission of the negative controls (substrate only), *Max* the average fluorescence emission from the positive controls (enzyme + substrate), and *Test* the emission from the assays in the presence of compounds (enzyme + compound + substrate). Dose-response confirmation assays were performed under similar conditions as the primary screen, but monitoring fluorescence at 10, 90, and 180 minutes. Tested concentrations of compounds were 2.5, 10, 20, 40, and 80 μM. The IC_50_ of each compound was determined using equation 2:

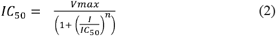

In the above, *n* measures cooperativity, with *n*>1 indicating synergistic binding of more than a single molecule of the compound.

### Parasite growth inhibition assay

The chloroquine resistant *P. falciparum* parasites (strain Dd2) were cultured in de-identified, IRB-exempt human red blood cells (RBCs) obtained from the American Red Cross and parasite cultures were maintained as described ^17^. Parasitaemia (percentage of total RBCs infected) was measured by flow cytometry on an Attune NxT (ThermoFisher) by nucleic-acid staining of cultured RBCs with phosphate-buffered saline (PBS) containing 0.8 □ μg □ ml^-1^ acridine orange. We began by determining percent growth inhibition at single compound dose (20 μM) for the 102 non-toxic compounds. Those that inhibited parasite growth by 50% or more were subjected to dose response analysis to extract EC_50_ values. The experiments were initiated with a starting parasitaemia of ∼1% and parasitaemia was measured 72 and 96 hours post compound addition. Growth was normalized to parasite cultures with carrier only (DMSO).

## Results and Discussion

### High-throughput screening of small molecule libraries to identify inhibitors of apPOL

To identify apPOL inhibitors we utilized our previously developed high-throughput DNA polymerase assay ^15^. The HT assay is based on a molecular beacon DNA template that forms a stem-loop with a Cy3 dye and the Black Hole Quencher on the 3′ and 5′ ends of the DNA, respectively. DNA polymerase activity extends an annealed primer through the stem region, displacing the fluorophore from the quencher, thereby increasing its fluorescence (Fig. 1A). The assay accurately recapitulates the kinetic behavior of apPOL, as measured through more conventional low-throughput gel-based assays ^18^. The Z′ factor for the assay described here was 0.9 across all plates (data not shown).

**Figure 1.**
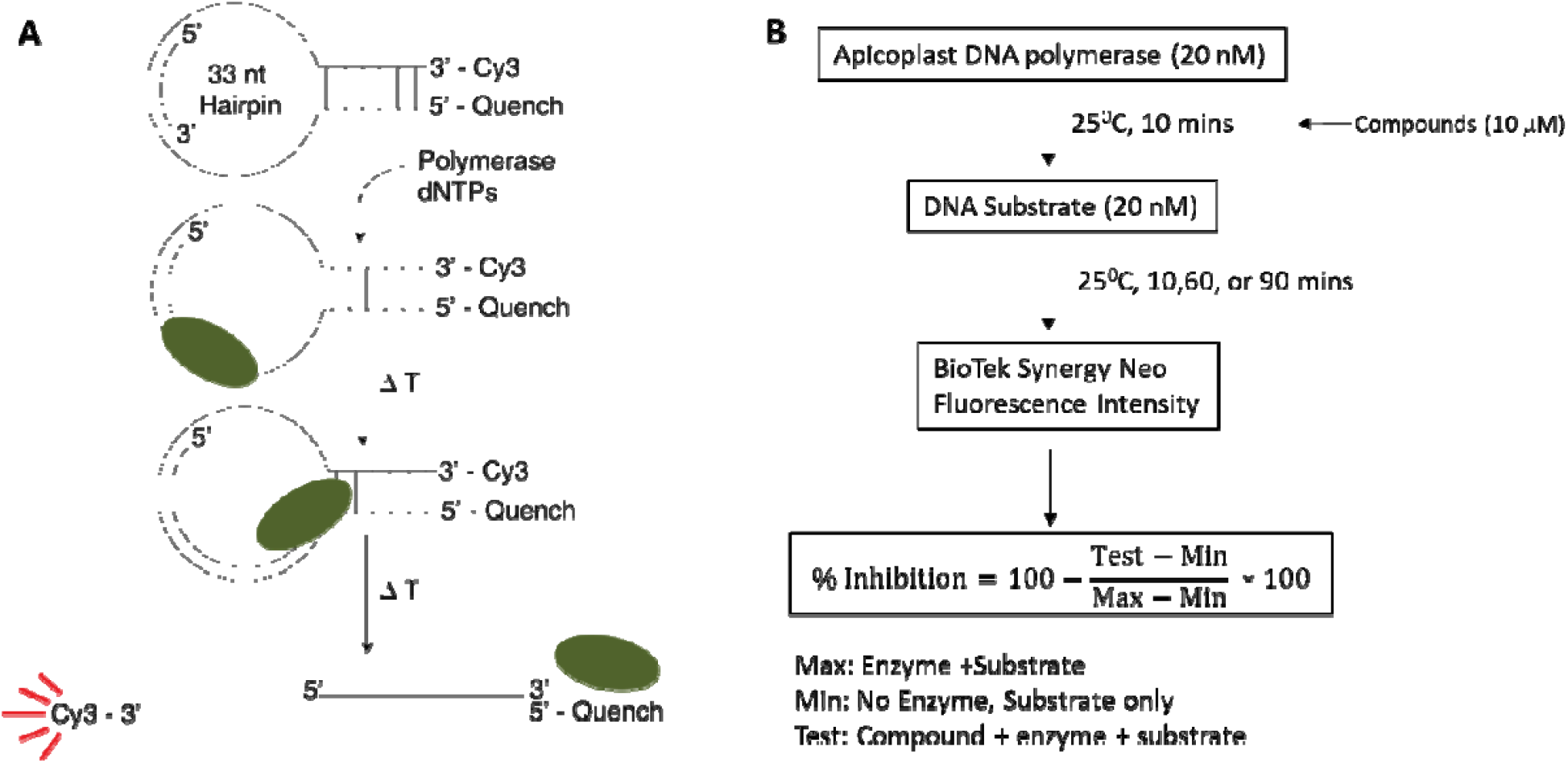
High-throughput screening. *A)* The fluorescent-based DNA polymerase assay use to screen the compound libraries. The DNA template strand is a hairpin loop with a primer annealed to the loop. The 3’ end of the primer is # bases away from the # bp duplex. Strand displacement activity of the DNA polymerase separates the Cy3 dye from the Black hole quencher leading to an increase in fluorescent signal. *B)* The high-throughput assay workflow. The DNA polymerase is incubated with 10 mM compound for 10 minutes prior to addition of the hairpin substrate. The reaction is quenched at 3 different time points and the fluorescent signal is measured and % inhibition is determined using the provided equation.

The initial screen followed the workflow outlined in Fig. 1B. The identity of the compound libraries screened and the resulting primary hits are given in Table 1 with a scattergram of the percent inhibition for each well shown in Fig. 2. A total of 206,504 compounds were tested for inhibition of *Pf*-apPOL with 1217 compounds qualifying as “hits” for an overall rate of 0.59%. Natural products and natural product-like compounds were overrepresented in the hit subgroup, with rates of 4.8% for the GreenPharma natural product library and 1.4% for the Life Chemicals natural product-like library. Also overrepresented (2.7%) was the Sigma Lopac 1280 library, which contains exclusively pharmacologically active compounds.

**Table 1.**
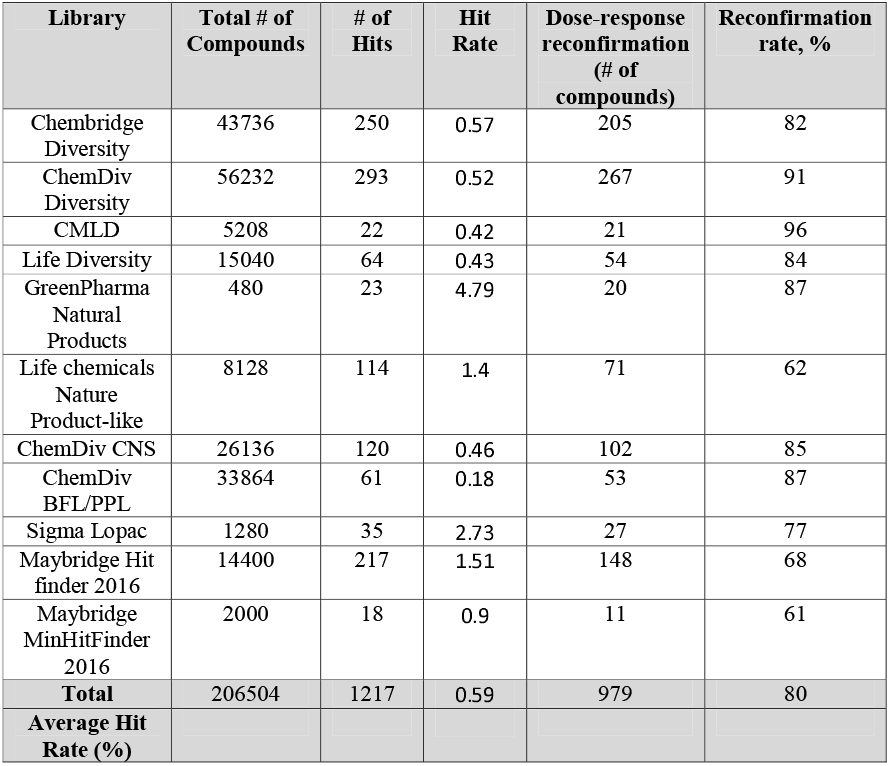
Results from high-throughput screening to identify inhibitors of *P. falciparum* apicoplast DNA polymerase.

**Figure 2.**
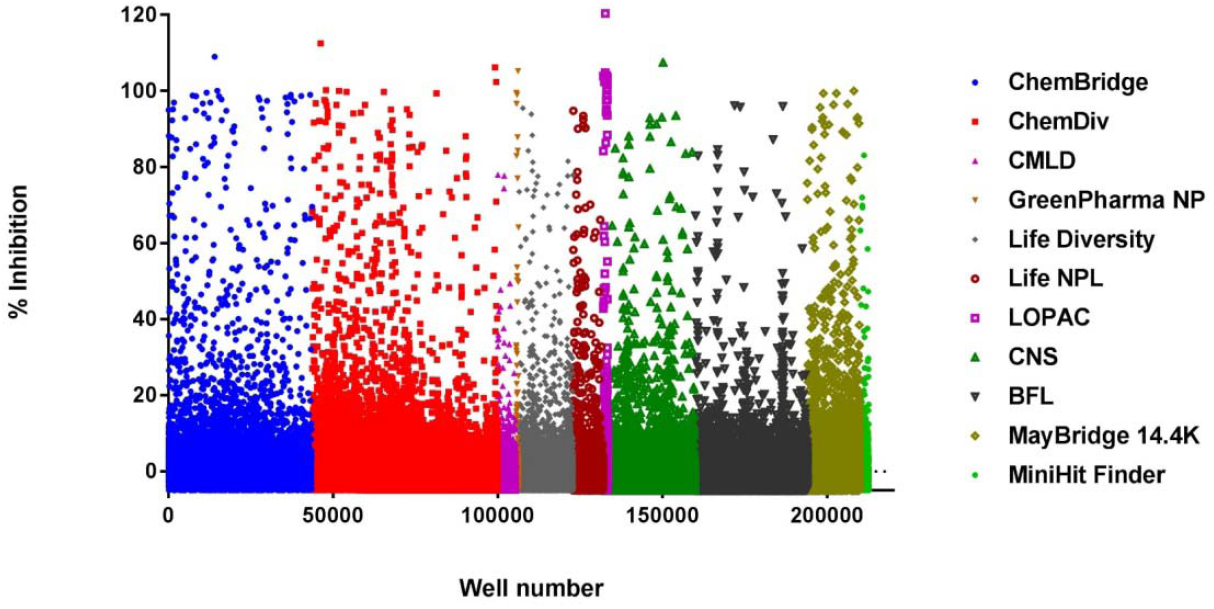
A scattergram of the large-scale screen of 11 compound libraries (listed at right). The overall hit rate for the screen was 0.59%

Each compound was tested in a dose-response format to determine IC_50_ values. Out of the 1217 compounds identified as hits, 979 exhibited potential inhibition in the dose-response reconfirmation assays (80.4%). These compounds were structurally diverse with only 8 clusters containing 10 or more compounds (supplemental Fig. S1). Example dose-response data for clusters with relatively high and low affinities are shown in supplemental Figures S2 and S3, respectively. The 979 likely to include compounds that interfere with the fluorescent read-out of the assay (false positives). (So a compound that does not inhibit but quenches Cy3 looks “active”. I think we should reserve terms like “active” to refer to enzyme activity to avoid literary contortions like “an active inhibitor inhibits enzyme activity”). Additionally, many of the identified compounds resemble DNA mimics or nucleotide analogs and are likely non-specific inhibitors of nucleic-acid polymerases and enzymes that bind nucleic acids. To identify compounds that specifically inhibit *P. falciparum* apPOL, we screened the 979 potential inhibitors against bacteriophage T4 DNA polymerase ^19^. T4 DNA polymerase can be produced in large quantities, and is in the same family (B-family) as the replicative human DNA polymerases (α, δ, and ε) 20. The majority of the 979 compounds (793) inhibited T4 DNA polymerase. Further investigation of these non-selective inhibitors of DNA polymerase could reveal modes of inhibition that are common across multiple DNA polymerase families. The remaining compounds (186) represent “primary” hits, that is, specific inhibitors of *P. falciparum* apicoplast DNA polymerase. On the basis of cluster analysis, primary hits are structurally diverse with only two clusters containing 10 or more compounds (supplemental Fig. S4). As shown in Fig. S4, the vast majority of the compounds (94%) do not violate any of the Lipinski’s rule of five (supplemental file 1).

### Toxicity of the primary hits toward human cells

As a measurement of toxicity, we employed the well-established CellTiter-Glo™ Luminescent Cell Viability Assay using HepaRG human cells. These cells contain endogenous luciferase enzyme that uses ATP and D-Luciferin to produce excited oxyluciferin, which then returns to its ground state by the emission of a photon (luminescence). Metabolically active cells generate ATP, hence luminescence is proportional to ATP present and an indication of cellular metabolic activity. The assay is robust, sensitive, reproducible, and routinely used in a high-throughput format at the KU-HTS facility ^21^. The results from the 48 hour cytotoxicity assay indicate that 102 of the 186 primary hits are non-toxic to the human hepatic cells (55%) at the highest concentration tested (100 μM). Cluster analysis indicates 68 groups of non-similar compounds, including clusters of 10 and 12 from the set of 186 apPOL-specific inhibitors (Fig. 3). In addition to analyzing the compounds for Lipinski violations, we also used SwissADME to determine the synthetic accessibility score and lead-likeness score for each compound ^22^. The synthetic accessibility score ranges from 1 (very easy) to 10 (very hard) and has been shown to correlate well with manual estimations of synthetic difficulty performed by chemists ^23^. The majority of the compounds appear to be synthetically tractable with scores ranging from 2-4 and only 7 molecules scoring above 4.0 (Fig. 4). Lead-likeness is determined through a combination of physiochemical filters (e.g, PAINS ^24^ and Brenk ^25^), and the notion that selection criteria for lead compounds should emphasize low molecular weights (100-350 Da) and degree of polarity (clgP) values of 1-3.0 because optimization of leads generally increase size and lipophilicity. As a consequence, leads are required to be smaller and less hydrophobic than drug-like molecules ^22,26^. From the set of 102 non-toxic apPOL inhibitors, 30 compounds have no lead-likeness violations and another 31 compounds only violate a single criteria, indicating that the majority of these compounds are potential leads.

**Figure 3.**
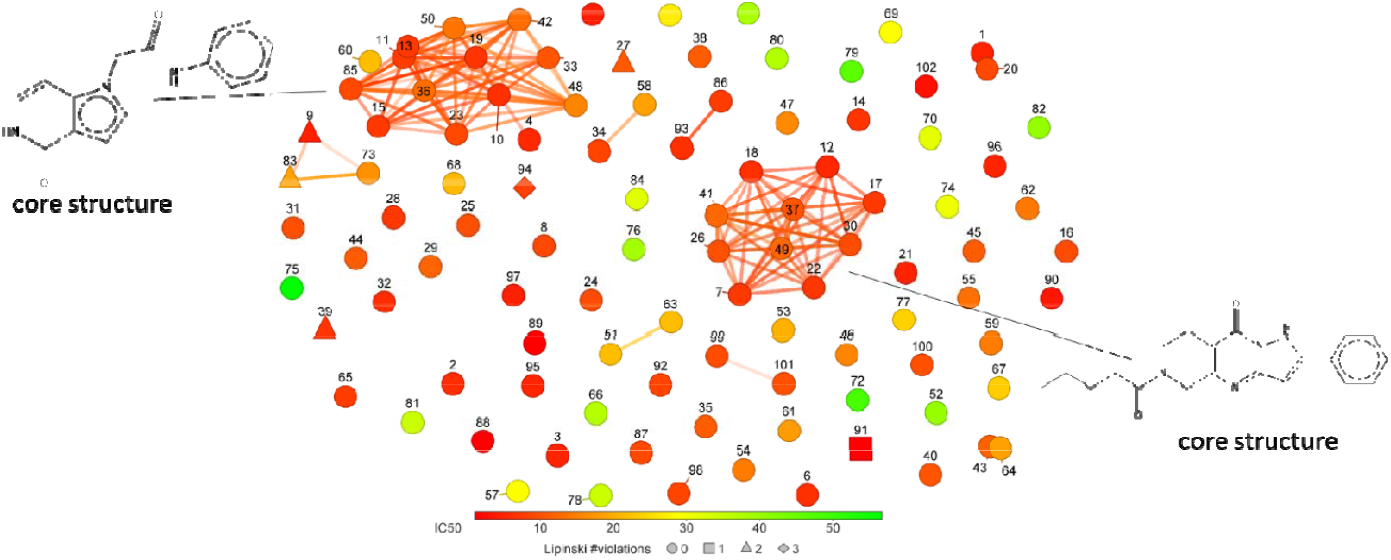
A plot of FragFp similarity for the nontoxic, specific apPOL inhibitors (102 compounds). Chemical similarity was calculated using the DataWarrior ^34^ software package. The core structure for the two largest clusters are given. Each compound is labeled with a number corresponding to the data in supplemental file 1. The markers are colored according to IC_50_ values and the number of Lipinski violations are indicated by marker shape.

**Figure 4.**
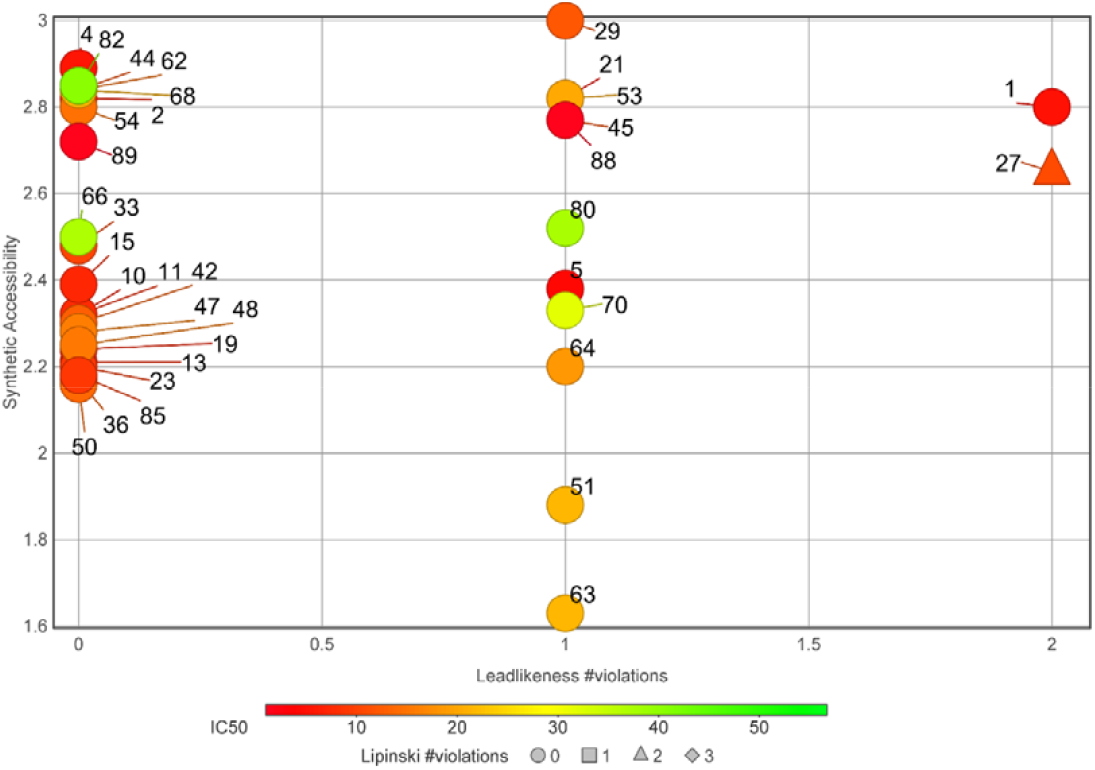
Potential as lead compounds. A plot of the synthetic accessibility *versus* leadlikeness violations for the non-toxic, specific apPOL inhibitors is shown. Each compound is labeled with a number corresponding to the data in supplemental file 1. The markers are colored according to their IC_50_ values and the number of Lipinski violations are indicated by marker shape. A description of synthetic accessibility and leadlikeness can be found in the main text.

### Lethality of compounds toward cultured P. falciparum parasites

We tested the ability of the 102 compounds to kill *P. falciparum* parasites cultured in human red blood cells. Due to the number of compounds, we used a single inhibitor concentration of 20 μM and performed the experiments in triplicate. These single-dose assays identified 9 compounds that inhibited parasite growth by more than 50%. These compounds were then tested in a dose-response format against chloroquine resistant *P. falciparum* parasites (Dd2) and the resulting data were fit to a sigmoidal growth inhibition curve to determine EC_50_ values (Figure 5, Table 2). Additionally, these compounds were tested for *in vitro* inhibition of *P. berghei* (*Pb*) and *Babesia bovis* (*Bb*) apicolast DNA polymerases, which are 82% and 43% identical to the *P. falciparum* protein, respectively. Inhibition of *Pb*-apPOL is a desired feature because it is likely that future *in vivo* studies will be performed in the mouse model for Malaria. Inhibition of *Bb-*apPOL, but not the more distantly related B-family polymerase T4 DNA pol, would suggest that the inhibitor may be pan-selective against most apicoplast apPOL’s. The *in vitro* polymerase assays identified 5 compounds that were potent inhibitors of all three apicoplast polymerases and a single compound that inhibited *Pf*-apPOL and *Bb*-apPOL but not *Pb*-apPOL (Table 2). Finally, to determine if the compounds were killing the parasites solely via apicoplast disruption ^27^, the dose-response curves were also performed with 1 mM isopentylpyrophosphate (IPP) in the growth media (Table 2). As seen in Table 2, none of the compounds display a significant IPP effect, suggesting that these compounds are hitting multiple cellular targets (e.g., apPOL and a non-apicoplast protein) or a specific non-apicoplast target. At this time, it is unclear what the non-apicoplast target(s) may be. One intriguing possibility is that these compounds may be targeting both the apicoplast and the mitochondrial DNA polymerases. Very little is known regarding mitochondrial DNA replication in *P. falciparum*; however, signal sequence identification algorithms have predicted that the other bacterial A-family DNA polymerase residing in *P. falciparum* is targeted to the mitochondria (PF3D7_0625300)^28^. As five of these *Pf*-apPOL inhibitors have the ability to inhibit the distantly related *Bb*-apPOL, it would not be surprising if they also inhibited the only other A-family polymerase in *P. falciparum*. The mitochondria has been recognized as a promising drug target ^29,30^ and saturating transposon mutagenesis in *P. falciparum*^31^ as well as pooled knockout screens in *P. berghei* ^32^ indicate that this A-family polymerase is likely essential. We are currently in the process of cloning and expressing this putative mitochondrial DNA polymerase to determine the effect of these compounds on its activity. If inhibition of the mitochondrial DNA polymerase is confirmed, these compounds would have very high potential for further development, since dual targeting both DNA polymerases residing within the parasite would increase potency and reduce the evolution of drug resistance.

**Table 2.** Inhibition data for compounds found to be active against cultured *Plasmodium falciparum*

| Molecule | KUID | IC<sub>50</sub><br>*Pf*-apPOL<br>(μM) | IC<sub>50</sub><br>*Pb*-apPOL<br>(μM) | IC<sub>50</sub><br>*Bb*-apPOL<br>(μM) | EC<sub>50</sub><br>(μM) | EC<sub>50</sub> + IPP<br>(μM) |
| --- | --- | --- | --- | --- | --- | --- |
| **1** | KU0263501 | 5.9 ± 0.9 | 7.1 ± 0.6 | 3.7 ± 0.2 | 8.6 ± 0.4 | 9.3 ± 0.5 |
| **21** | KU0036696 | 8 ± 1 | 9.7 ± 1.6 | 8 ± 2 | 3.1 ± 0.1 | 3.4 ± 0.1 |
| **27** | KU0260920 | 15 ± 1 | ND | ND | 10 ± 1 | 9 ± 1 |
| **31** | KU0007309 | 15 ± 1 | ND | 15 ± 5 | 8.9 ± 0.5 | 8.7 ± 0.6 |
| **34** | KU0241474 | 7.2 ± 0.9 | ND | ND | 11.3 ± 0.9 | 10.0 ± 0.7 |
| **55** | KU0177470 | 10 ± 2 | 51.9 ± 3.5 | 1.5 ± 1.0 | 3.5 ± 0.3 | 4.3 ± 0.2 |
| **94** | KU0261556 | 7.6 ± 0.6 | 13.9 ± 1.6 | 30 ± 3 | 4.6 ± 0.2 | 3.7 ± 0.5 |
| **95** | KU0001071 | 7.4 ± 0.4 | 8.6 ± 0.7 | 2.5 ± 0.2 | 7.5 ± 0.5 | 7.4 ± 0.4 |
| **98** | KU0271653 | 8 ± 2 | ND | ND | 16 ± 1 | 11 ± 1 |

**Figure 5.**
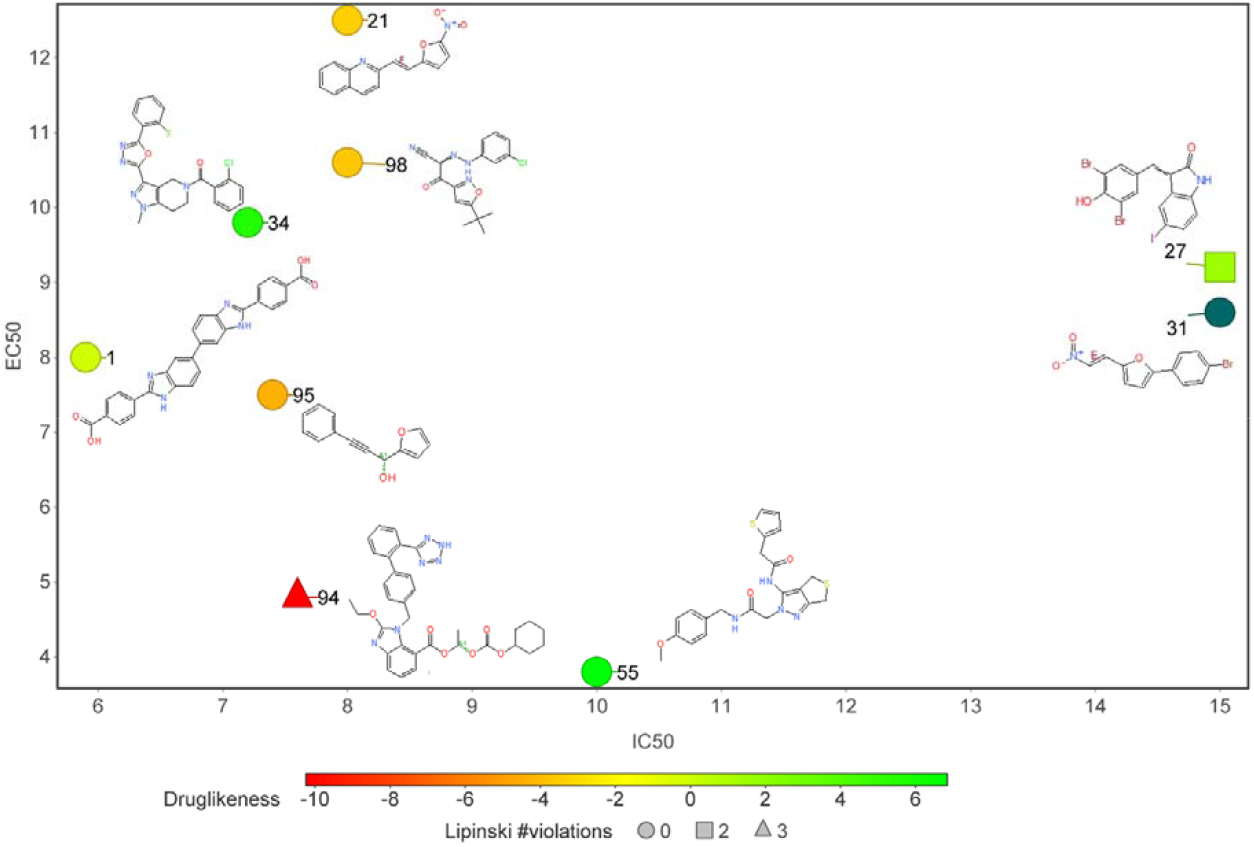
Compounds active against cultured *Plasmodium falciparum*. A plot of IC_50_ values versus EC_50_ values is shown. The markers are colored according to their Druglinkess values as calculated by the DataWarrior ^34^ software package. The number of Lipinski violations are indicated by marker shape.

## Conclusions

Resistance to current Malaria drugs is inevitable and new drugs that target novel aspects of *P. falciparum’s* biology are necessary to combat the disease. The apicoplast is a validated drug target and inhibition of apicoplast DNA replication leads to the loss of the organelle and parasite death^33^. The large-scale high-throughput screen described here defined several sets of compounds that could serve as tools to investigate mechanisms of DNA polymerase inhibition or as lead compounds for further optimization into anti-malaria drugs. Future work will involve determining mechanisms of inhibition, apPOL binding sites, and target(s) confirmation.

## Figure Legends

**Figure S1.**
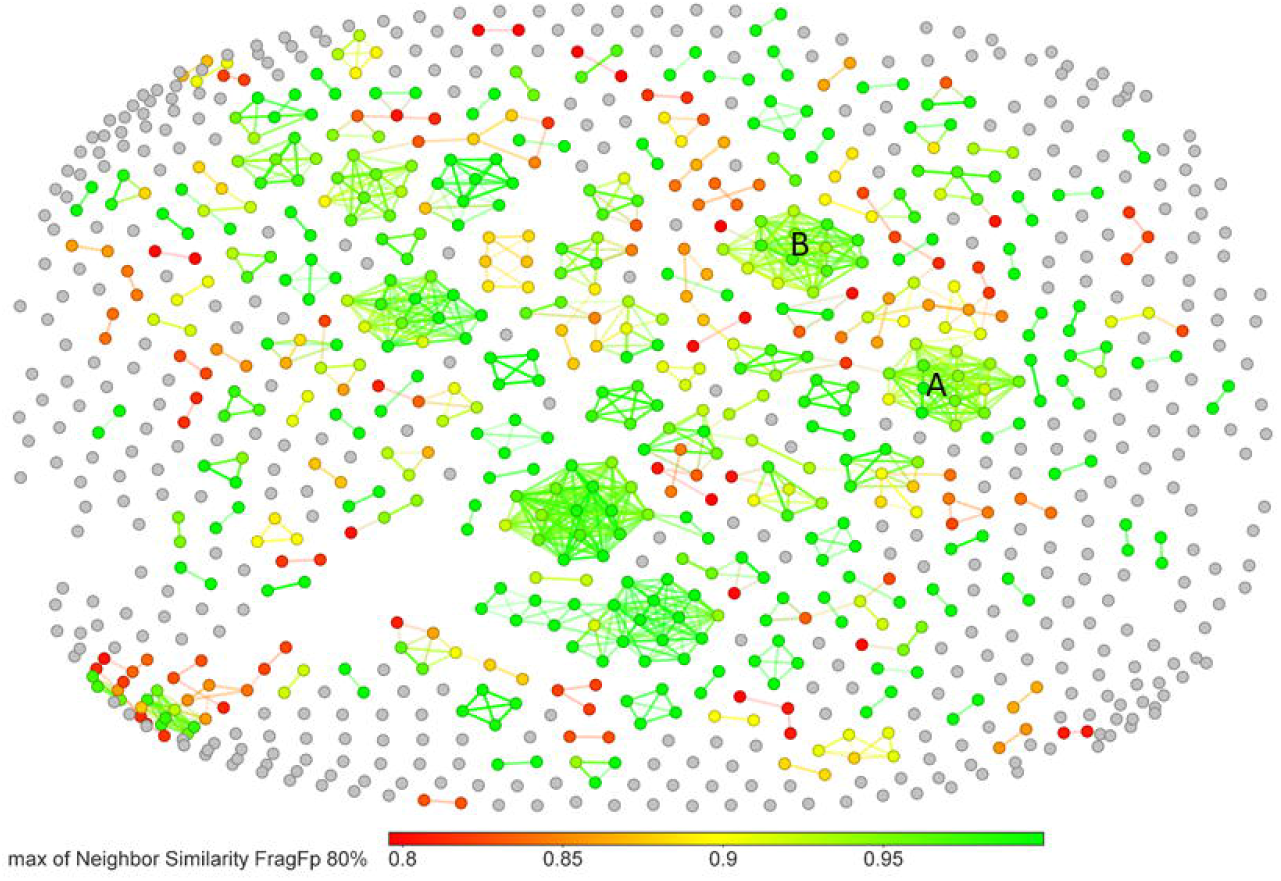
A plot of FragFp similarity for all validated primary hits (979 compounds). Chemical similarity was calculated using the DataWarrior^ref^ software package. The two largest clusters are indicated by the letters A and B and their structures and dose-response curves can be found in Figures S2 and S3.

**Figure S2.**
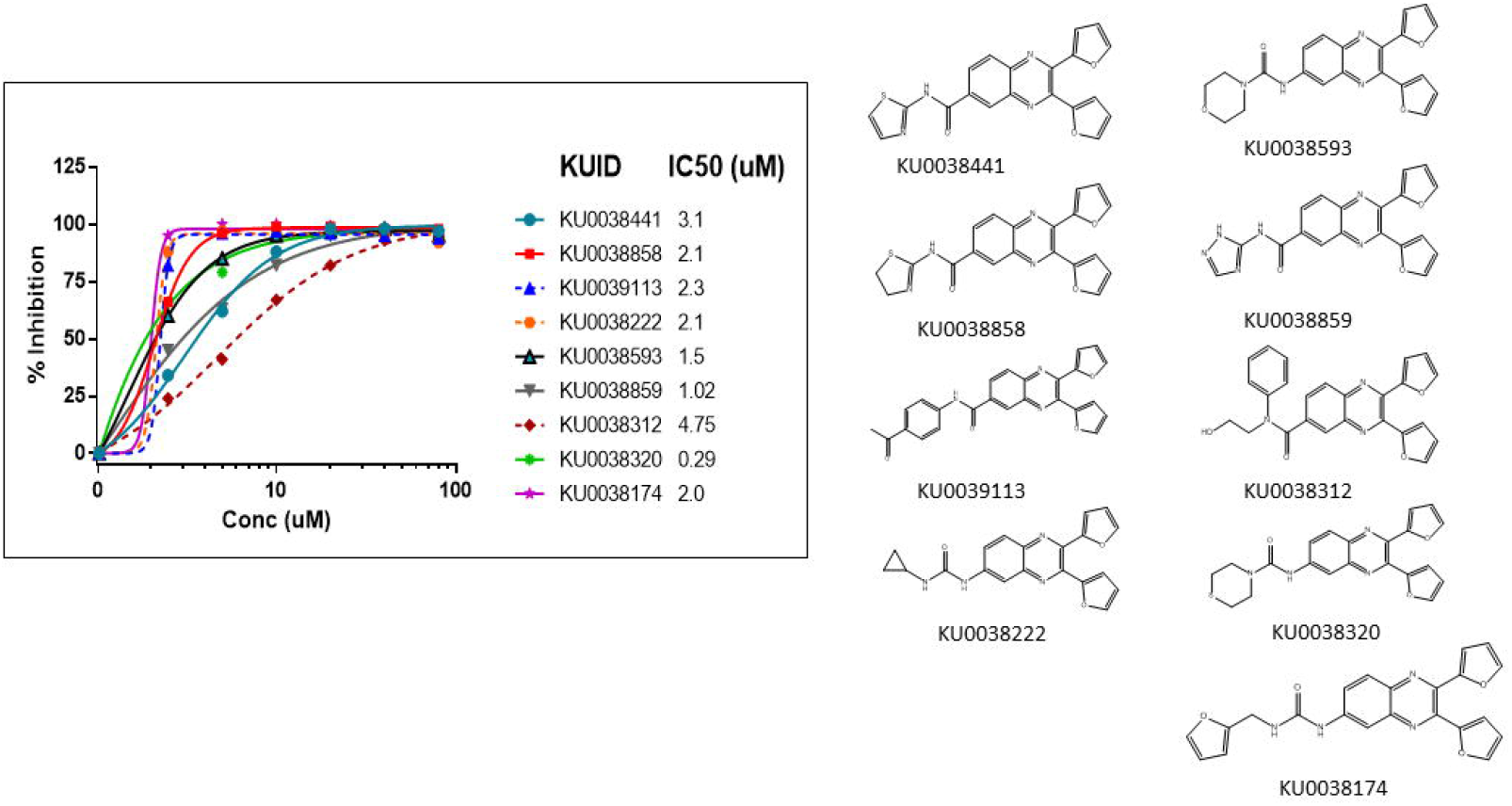
Chemical structures and dose-response data and for the compounds found in Cluster A from Figure S1.

**Supplementary Figure S3.**
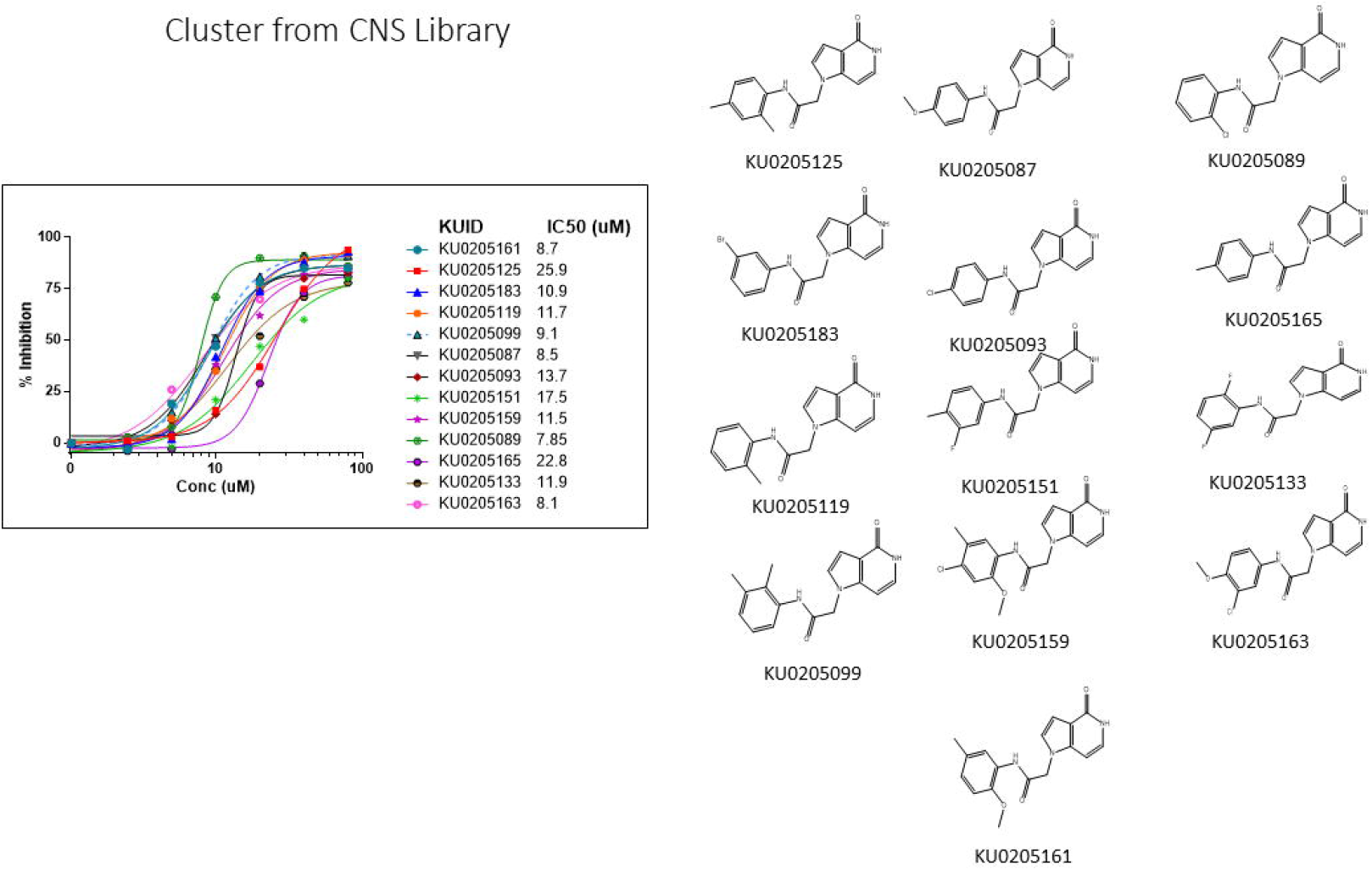
Chemical structures and dose-response data and for the compounds found in Cluster B from Figure S1.

**Figure S4.**
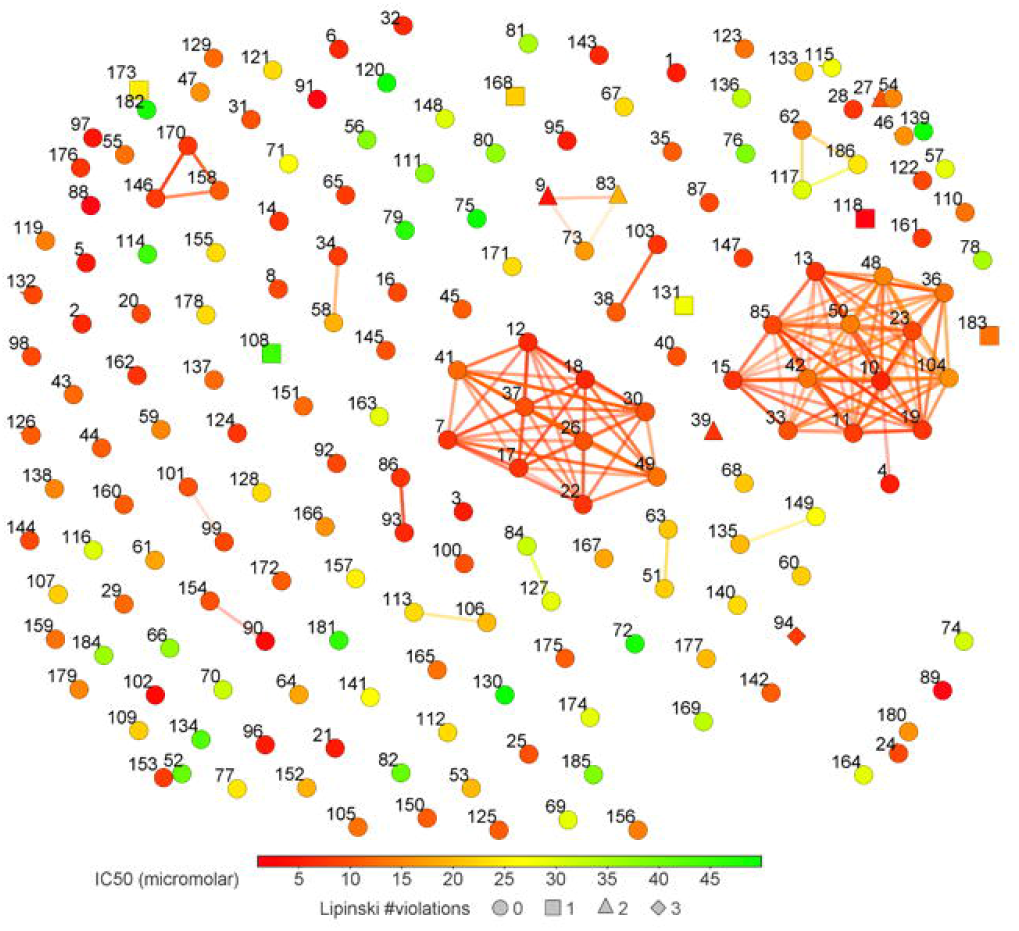
A plot of FragFp similarity for the specific apPOL inhibitors (183 compounds). Chemical similarity was calculated using the DataWarrior^ref^ software package. Each compound is labeled with a number corresponding to the data in supplemental file 1. The markers are colored according to IC_50_ values and the number of Lipinski violations are indicated by marker shape.

